# The illusion of interpretability in biologically informed neural networks

**DOI:** 10.64898/2026.05.07.723544

**Authors:** Isabella Caranzano, Tiziana Sanavia, Pietro Liò, Pierre Baldi, Piero Fariselli

**Affiliations:** AI and Computational Biomedicine Unit, Department of Medical Sciences, University of Turin, via Santena 19, 10126 Turin, Italy; Department of Computer Science and Technology, University of Cambridge, Cambridge, UK; Department of Computer Science, University of California, Irvine, CA 92697, USA

## Abstract

Biologically informed neural networks (BINNs), also known as visible neural networks (VNNs), are widely adopted in omics because their architectures mirror known biological structures, such as gene-to-pathway relationships, and are therefore often assumed to be inherently interpretable. This assumption implies that learned gene-to-pathway weights and pathway node activations reflect meaningful biological mechanisms. Here, we show that this premise fails for a classical reason: nonidentifiability. Using a controlled teacher and student framework, we demonstrate that even under ideal conditions, including noiseless data, the correct model class, and identical sparse wiring, a BINN can perfectly recover the input-to-output mapping while failing to recover both gene-to-pathway weights and pathway activations. This failure persists across classification, regression, and survival tasks, and remains robust to variations in biological structure and network depth. Thus, the problem is not merely overparameterization or poor optimization: learning from outputs alone does not identify internal structure. Since biological mechanisms are not directly observed, recovering them from predictions alone is *harder, not easier*, than recovering neural network parameters, which are already known to be nonidentifiable. Critically, this failure reflects standard practice: widely used BINNs do not impose objective level constraints on gene-to-pathway weights or pathway activations, and therefore operate precisely in the regime modeled by our teacher-student framework. These results indicate that architectural transparency does not imply mechanistic interpretability. Without constraints that explicitly enforce identifiability, the apparent interpretability of BINNs reflects their design rather than what they actually learn.

## 1 Introduction

Biologically informed neural networks (BINNs) are an excellent starting point for integrating curated biological knowledge into predictive modeling [1]. Their appeal is clear: genes feed into pathway modules or other interaction networks, which then feed into phenotypes, providing an intuitive *mechanistic map* that is easy to communicate and interpret. BINNs are also referred to as visible neural networks (VNNs), since the goal is to attribute semantic information to internal nodes [2]. In practice, BINNs have been applied to a broad range of omics prediction tasks, including regression, classification, and survival modeling [2].

Here, we focus specifically on BINNs and VNNs whose topology is constrained by prior biological knowledge, encoded through the wiring diagram of the network, and in which interpretability is expected to arise directly from this structure [3–20]. This is the setting in which interpretability is most directly assumed: genes are connected to pathway nodes, pathway nodes are labelled with biological meaning, and learned weights or node activations are often interpreted as biological quantities. Importantly, interpretability in these models is not enforced through objective-level constraints, such as sign regularization, monotonicity penalties, or experimental anchoring of pathway weights, but is instead inferred directly from architectural structure alone. The regime considered here should therefore be distinguished from physics informed neural networks [21] and biology informed neural networks in the sense of BioPINNs [22], where prior knowledge is imposed through explicit equations or constraints in the loss function. Our analysis targets topology based BINNs and VNNs, which are widely used in omics precisely because their architecture appears directly interpretable.

The basic idea is to obtain explanations that are not only based on post hoc interpretability tools, such as saliency maps or SHAP values [23], but are encoded directly in the model structure. In this setting, interpretability is typically understood at two complementary levels. At the parameter level, gene-to-pathway weights are interpreted as biologically meaningful associations. At the representation level, the activations of pathway nodes are often treated as proxies for pathway activity, and therefore as biologically informative intermediate features.

We raise a narrow but fundamental point: *the wiring diagram alone is generally not sufficient to ensure that the learned internal connections or activations correspond to biological relationships*. This is not a new pathology of neural networks, but an instance of a classical problem, well known in machine learning: the lack of identifiability. Distinct internal parameterizations may implement the same input-to-output function, so agreement at the output level does not imply recovery of internal weights or representations; see, for instance, [24–26].

This point is especially important for BINNs. In ordinary neural networks, even the recovery of internal weights from input-to-output observations is not guaranteed. In biological applications, the target is even more ambitious: one aims to recover biological mechanisms that are not directly observed, using noisy and partial omics measurements. Thus, if internal structure is not identifiable even in an idealized teacher-student setting with noiseless data, correct model class, and known sparse wiring, *it cannot be assumed to be identifiable in real biological applications without additional constraints*.

To isolate the fundamental limitations of this assumption, we adopt a controlled teacher-student framework. A teacher network defines a ground truth gene-to-pathway mapping and generates synthetic data, while a student network with identical architecture and wiring is trained to reproduce the teacher’s outputs. This setting represents a best case scenario in which the model class is correct, the data are noiseless, and the true biological structure is known. Under these conditions, one would expect both gene-to-pathway weights and pathway level representations to be recoverable.

As we show, this is not the case. Even in this idealized setting, learning the correct input-to-output mapping does not imply recovery of either the underlying biological parameters or the corresponding pathway activations. This indicates that interpretability fails not only at the level of weights, but also at the level of internal representations, and suggests that additional constraints are required to meaningfully anchor these models to biological mechanisms.

Although we focus on gene-to-pathway relationships in omics data as a concrete case study, the conclusions are not specific to this biological setting. The underlying issue is structural: it arises from the use of topology constrained architectures trained with output only objectives. As a result, the lack of identifiability demonstrated here applies generally to biologically informed and visible neural networks, independently of the specific entities, such as genes, pathways, proteins, or other biological modules, used to define the network topology.

## 2 Methods

### 2.1 Rationale and computational framework

Biological regulation is complex, and in real omics applications we only observe partial, noisy, and context dependent measurements of the underlying mechanisms. To isolate the fundamental identifiability problem from these biological and experimental confounders, we adopted a *deliberately simplified* and *favorable* setting. We assumed that, for the tasks considered here, the relevant biology could be approximated by a neural network in which gene level inputs are first aggregated into pathway level nodes through a sparse biologically informed layer. This abstraction captures the core architectural idea behind biologically informed neural networks (BINNs) and visible neural networks (VNNs): restricting connections according to prior biological knowledge in order to promote interpretability.

All experiments were conducted within a controlled *teacher and student* framework. A teacher neural network (TNN, Fig. 1 panel **a**) was first defined to represent the ground truth input-to-output mapping, corresponding to the unknown biological mechanism in this idealized setting. A student neural network (SNN, Fig. 1 panel **b**), sharing the same architectural class and sparse wiring diagram, was then trained from random initialization, exclusively through knowledge distillation, to match the teacher’s outputs.

**Figure 1:**
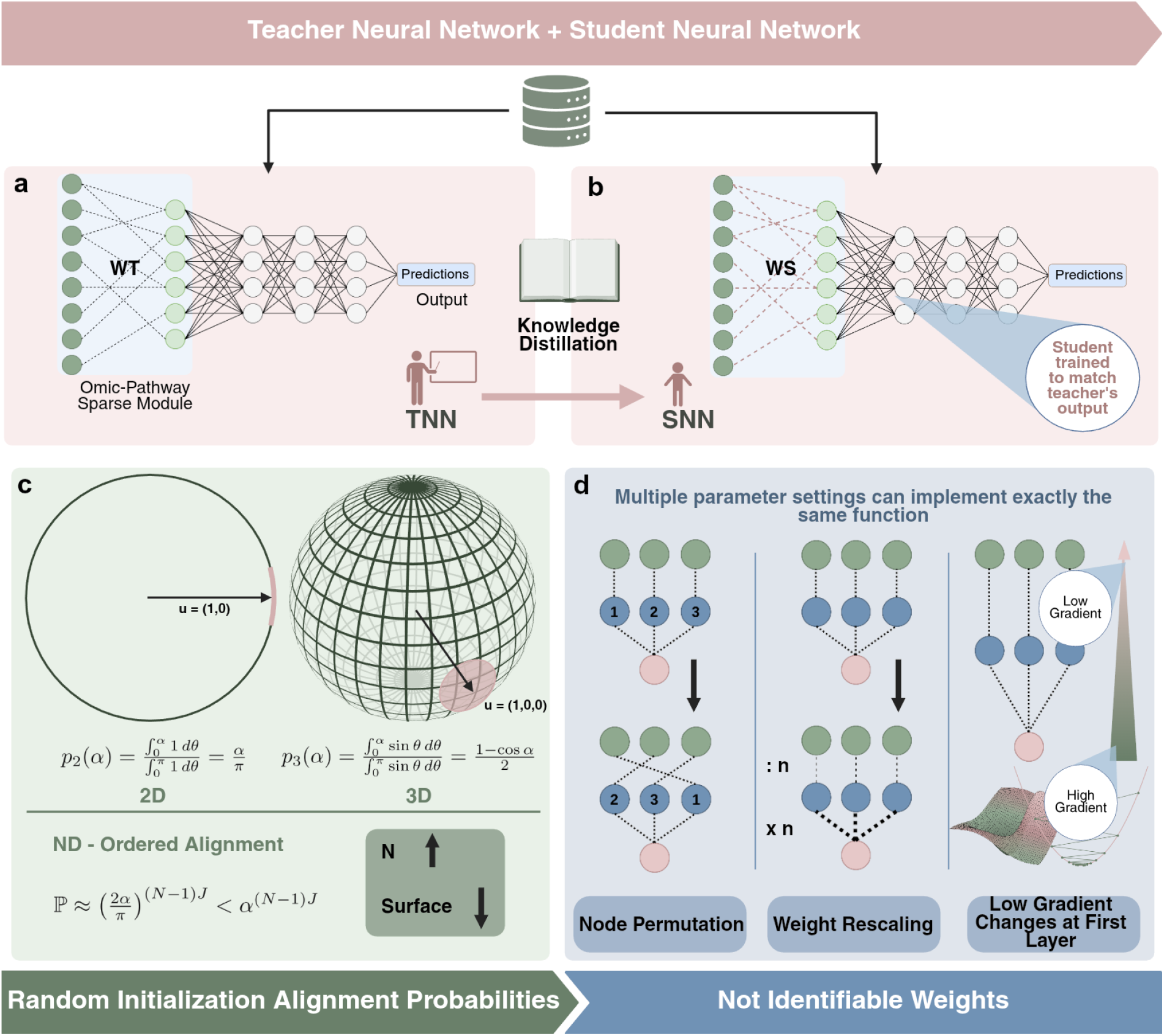
Teacher–student analysis reveals fundamental limits of pathway interpretability in biologically-informed neural networks. (**a**) The Teacher neural network (TNN, weights indicated by WT) implementing a “hypothetically true” biologically structured gene–pathway mapping, followed by a dense neural backbone, used to generate predictions from simulated omics data. (**b**) Student neural network (SNN, weights indicated by WS) with identical architecture and sparse wiring, trained via knowledge distillation to match the teacher’s output, without explicit constraints on the internal pathway representation. (**c**) Geometric intuition for random initialization in high-dimensional spaces: the probability that a randomly initialized pathway weight vector is positively aligned with a fixed ground-truth direction decays rapidly as dimensionality increases, making correct alignment unlikely for large pathway sizes.(d)Schematic illustration of nonidentifiability mechanisms in deep networks, including node permutation, weight rescaling with downstream compensation, and weak or non-directional gradient signals reaching the first layer, all of which allow multiple distinct parameter configurations to implement the same input–output function. Together, these effects explain how biologically-informed architectures can achieve accurate predictions while failing to recover biologically meaningful first-layer representations.

Importantly, the student received no direct supervision, regularization, or alignment constraint on internal weights or pathway activations. The training objective constrained only the network output. This design allowed us to test whether accurate functional matching is sufficient to recover biologically meaningful internal structure, both at the level of gene-to-pathway weights and at the level of pathway node activations.

The mathematical formulation of this problem, including the analysis of initialization misalignment and lack of identifiability of first layer weights, is provided in detail in the Supplementary Material, so that the argument can be read independently. Related forms of this problem have also been discussed in the machine learning literature (see, for instance, [24–26]). Briefly, the Supplementary analysis shows that, for a pathway node with *s* incoming genes, the probability that a random isotropic initialization is aligned with a fixed ground truth weight vector decays rapidly with *s*. This directional analysis is a simplifying restriction, since it ignores weight magnitudes and considers only alignment. When weight magnitudes are also considered, as in the simulations, the space of possible solutions is even larger and recovery becomes harder. The Supplementary Material also shows that, even under perfect prediction, multiple distinct internal parameterizations can implement exactly the same function, due to mechanisms such as node permutation symmetries, weight rescaling invariances across layers, and weak or distorted gradients reaching the first biologically interpreted layer.

### 2.2 Simulated data generation

To eliminate confounding effects due to unknown biology, measurement noise, or model misspecification, all experiments were performed on simulated data generated from a known teacher model. In the reference configuration, input samples consisted of *G* = 400 gene-level features. Pathway structure was defined by *P* = 60 pathway nodes, each connected to a subset of genes with pathway sizes uniformly sampled between 14 and 22 genes. Pathways were allowed to overlap, with an average overlap fraction of 0.55, reflecting the fact that genes often participate in multiple biological processes.

Training and test datasets comprised 12,000 and 3,000 samples, respectively, drawn independently using fixed random seeds to ensure reproducibility. Input features were sampled from a standard multivariate distribution and propagated through the teacher network to generate task-specific outputs. For survival analysis, event times were generated from teacher-defined risk scores using a proportional hazards model, with an independent censoring rate of 5%.

This simulation design represents *an idealized best-case scenario for interpretability* : the true data-generating process is exactly representable by the model class, the biological mask is known, and the data are noiseless. Therefore, any failure to recover the teacher’s internal weights or pathway activations cannot be attributed to incomplete biology or noisy measurements, but instead reflects limitations arising from optimization and identifiability.

### 2.3 Network architecture

Both teacher and student networks followed the same modular architecture, composed of three conceptual blocks: a biologically informed gene-pathway layer, a dense neural backbone, and a task-specific output layer.

#### Gene-pathway layer

The first layer mapped gene-level inputs to pathway-level representations through a fixed binary mask

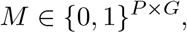

encoding gene membership within pathways. Each pathway node received input only from its associated genes, and all connections outside the mask were fixed to zero throughout training. Within the mask, weights were trainable and unconstrained. No normalization, sign constraints, weight tying, or pathway-level regularization was applied unless explicitly stated.

This layer represents the primary source of interpretability in topology-based BINNs and VNNs, because pathway nodes are explicitly labelled and are intended to correspond to biological entities.

#### Dense backbone

The pathway layer was followed by one or more fully connected hidden layers forming a dense neural backbone. Each hidden layer applied an affine transformation followed by a ReLU nonlinearity. These layers were unconstrained and allowed to freely mix pathway-level signals. The backbone provided sufficient expressive capacity to compensate for imperfect or non-biological representations learned in the first layer.

#### Output layer

The final layer was task-dependent and mapped the last hidden representation to the predicted output. No parameter sharing or additional alignment constraint was imposed between teacher and student beyond architectural equivalence and the shared sparse mask.

### 2.4 Prediction tasks

We evaluated the teacher-student framework across four representative omics prediction settings: binary classification, multiclass classification, regression, and survival analysis. In all cases, the student was trained by knowledge distillation to reproduce the teacher’s outputs rather than external ground-truth labels. Predictive performance was assessed using task-appropriate metrics, including classification accuracy, mean squared error, coefficient of determination *R*^2^, and concordance index for survival analysis. Full task-specific loss functions and evaluation details are reported in the Additional Information.

### 2.5 Training protocol and repetitions

Student networks were trained using gradient-based optimization until convergence of the distillation objective, as verified by near-zero matching loss and stable predictive performance. Teacher parameters were fixed throughout student training.

For each task, 20 independent student instances were trained from different random initializations. This repetition enabled us to quantify the variability of internal solutions across functionally equivalent models and to assess whether recovery of the teacher’s biological representation was robust to initialization.

Unless otherwise specified, models were trained using a batch size of 512, a learning rate of 2 × 10^−3^, and 120 training epochs. No weight decay was applied. Dropout was used only in the teacher network, with dropout rate 0.15, to avoid trivial memorization, whereas student networks were trained without dropout.

### 2.6 Depth sensitivity analysis

To determine whether nonidentifiability was primarily a consequence of deep overparameterized architectures, we systematically varied the depth of the dense backbone. Starting from a shallow model with a single hidden layer, we progressively increased network depth up to 25 hidden layers, resulting in 25 depth configurations. Each hidden layer contained 64 neurons and used ReLU activations, while the gene-pathway mask, data-generating process, loss functions, and optimization protocol were kept fixed.

This analysis was designed to test whether recovery failure occurs only in deep networks, where downstream layers have greater capacity to compensate for incorrect early representations, or whether it is already present in shallow architectures. Although deeper models are expected to exacerbate nonidentifiability, the key question was whether shallow models are sufficient to restore biologically meaningful recovery of first-layer weights and pathway activations.

### 2.7 Structural sensitivity analysis of the simulated biology

To verify that our conclusions did not depend on one arbitrary choice of simulated biological structure, we performed additional experiments varying the main structural parameters of the gene-pathway system. Specifically, we varied the number of genes, the number of pathways, the degree of overlap between pathways, and the distribution of pathway sizes. These analyses were designed to test whether the failure of internal recovery was specific to the reference configuration, or whether it persisted across different plausible biological organizations.

For each structural configuration, the teacher and student shared the same sparse mask and architectural class, and the student was again trained only to match the teacher’s outputs. Recovery was then evaluated using the same parameter-level and activation-level metrics described below.

### 2.8 Evaluation of internal recovery

Internal recovery was evaluated at two complementary levels: parameter-level interpretability and representation-level interpretability. For parameter-level recovery, we compared corresponding teacher and student gene-pathway weight vectors using relative *L*_2_ error and cosine similarity. For representation-level recovery, we compared teacher and student pathway-node activations on held-out samples using Pearson correlation, sample-wise cosine similarity, and the coefficient of determination *R*^2^. Full definitions of these metrics and implementation details are provided in the Additional Information.

### 2.9 Implementation details

All experiments were implemented in Python using PyTorch and run on an NVIDIA RTX 4070 GPU. Random seeds were controlled at initialization to ensure reproducibility. Training was performed on minibatches using standard gradient-based optimizers. All results were automatically logged and stored for downstream statistical analysis.

## 3 Results

Across all tasks, student networks achieved near-perfect agreement with the teacher’s predictions, confirming that the correct input-to-output mapping was successfully learned. In classification settings, accuracy approached 1.0, while regression and survival tasks achieved high coefficients of determination (*R*^2^ ∼ 0.73) and concordance indices (C-index ∼ 0.89), respectively (Fig. 2 panel **b**). However, this functional agreement did not extend to the internal structure of the network.

**Figure 2:**
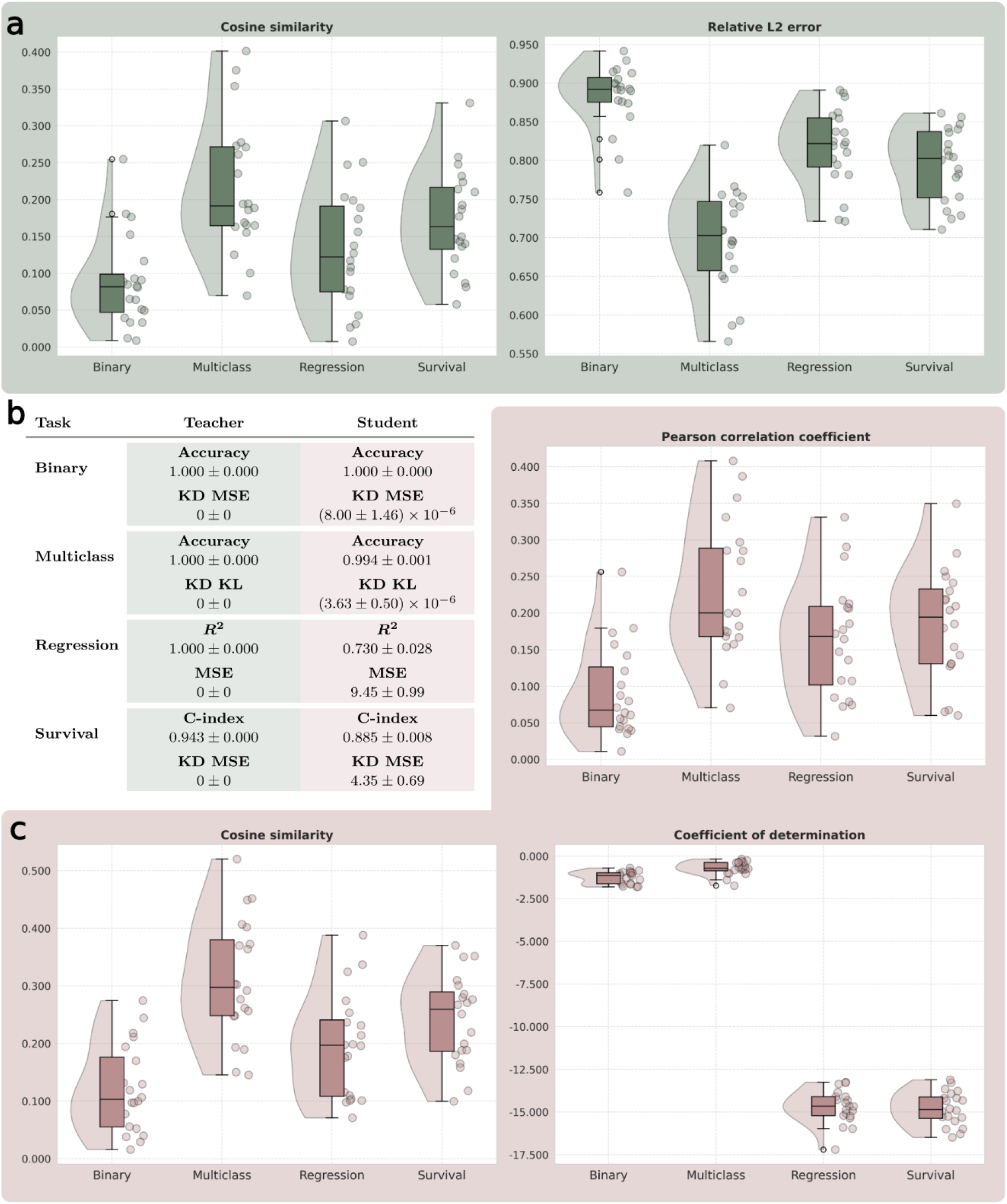
Failure of interpretability despite accurate functional learning. (**a**) **Parameter-level interpretability**. Cosine similarity and relative *L*_2_ error between teacher and student gene–pathway weights across binary classification, multiclass classification, regression, and survival tasks. Despite identical architectures and near-perfect output matching, student networks fail to recover the teacher’s first-layer parameters, showing low angular alignment and high reconstruction error. (**b**) **Teacher-student predictive performance**. Comparison of teacher and student models across the four tasks. Predictive performance is nearly identical, confirming successful functional learning despite discrepancies in internal representations. (**c**) **Activation-level interpretability**. Pearson correlation coefficient and coefficient of determination (*R*^2^) between teacher and student pathway-node activations. Low correlation and poor *R*^2^ values indicate that pathway-level representations are not preserved, demonstrating that interpretability fails also at the level of node activations.

At the parameter level, the student consistently failed to recover the teacher’s gene-to-pathway weights. Relative errors remained high, typically between 0.7 and 0.9, and cosine similarity remained low, with median values in the range 0.05 to 0.20. This indicates that the learned weights were largely misaligned with the biological associations encoded in the teacher model (Fig. 2 panel **a**). This failure occurred despite the fact that the student architecture exactly matched the data generating process, and that the training objective was fully consistent with the teacher’s outputs. In biological terms, this means that even when the phenotype is predicted almost perfectly, and even when the biological wiring is assumed to be correct, for example the links between genes and pathways or between proteins, the inferred strengths of those links can still be wrong.

Crucially, this mismatch also persisted at the level of internal representations. Pathway node activations in student networks were only weakly aligned with those of the teacher, with median correlations typically in the range 0.05 to 0.20 and low cosine similarity. Strikingly, the coefficient of determination between teacher and student activations was frequently negative, in some cases with *R*^2^ *<* −10, indicating that student representations reconstructed the teacher’s pathway level signals worse than predicting the mean teacher activation (Fig. 2 panel **c**). Thus, even when predictions were indistinguishable, the internal biological representations were not preserved.

These results show that assigning a biological name to an internal node does not by itself make the learned activation of that node a biologically meaningful quantity. If a student network with the same inputs, outputs, architecture, and wiring can reproduce the teacher perfectly while learning different weights and activations, then the semantic label attached to the node is an architectural annotation, not an identified biological variable. In machine learning terms, the student learns the same function but not the same internal parameterization or representation. In biological terms, the model can reproduce the phenotype while failing to recover the intended gene-to-pathway weights or pathway activities. Therefore, interpretability in biologically informed neural networks cannot be assumed at either the parameter or representation level.

To assess the robustness of the observed nonidentifiability beyond a single reference configuration, we performed additional analyses by systematically varying key structural properties of the simulated biological system as well as the depth of the network architecture.

Across all structural perturbations, recovery of gene-pathway weights remains consistently poor. Systematic changes in the number of genes, number of pathways, pathway sizes, and degree of overlap did not qualitatively alter the results. Across all configurations, high predictive agreement consistently coexisted with poor recovery of both weights and activations, indicating that the failure to recover meaningful internal structure is a general property of the model class (Fig. 3). Increasing the number of pathways or the size of pathways systematically worsens alignment, reflecting the expansion of the space of functionally equivalent internal solutions. Similarly, increasing the number of genes does not lead to meaningful improvements, with recovery rapidly saturating at high error levels. Introducing biologically realistic features such as pathway overlap does not resolve this issue and, in some cases, further degrades alignment, likely due to increased redundancy and ambiguity in pathway representations. Taken together, these results indicate that the failure to recover biologically meaningful parameters is not tied to a specific choice of simulated biology, but instead reflects a general structural nonidentifiability inherent to topology-constrained BINNs.

**Figure 3:**
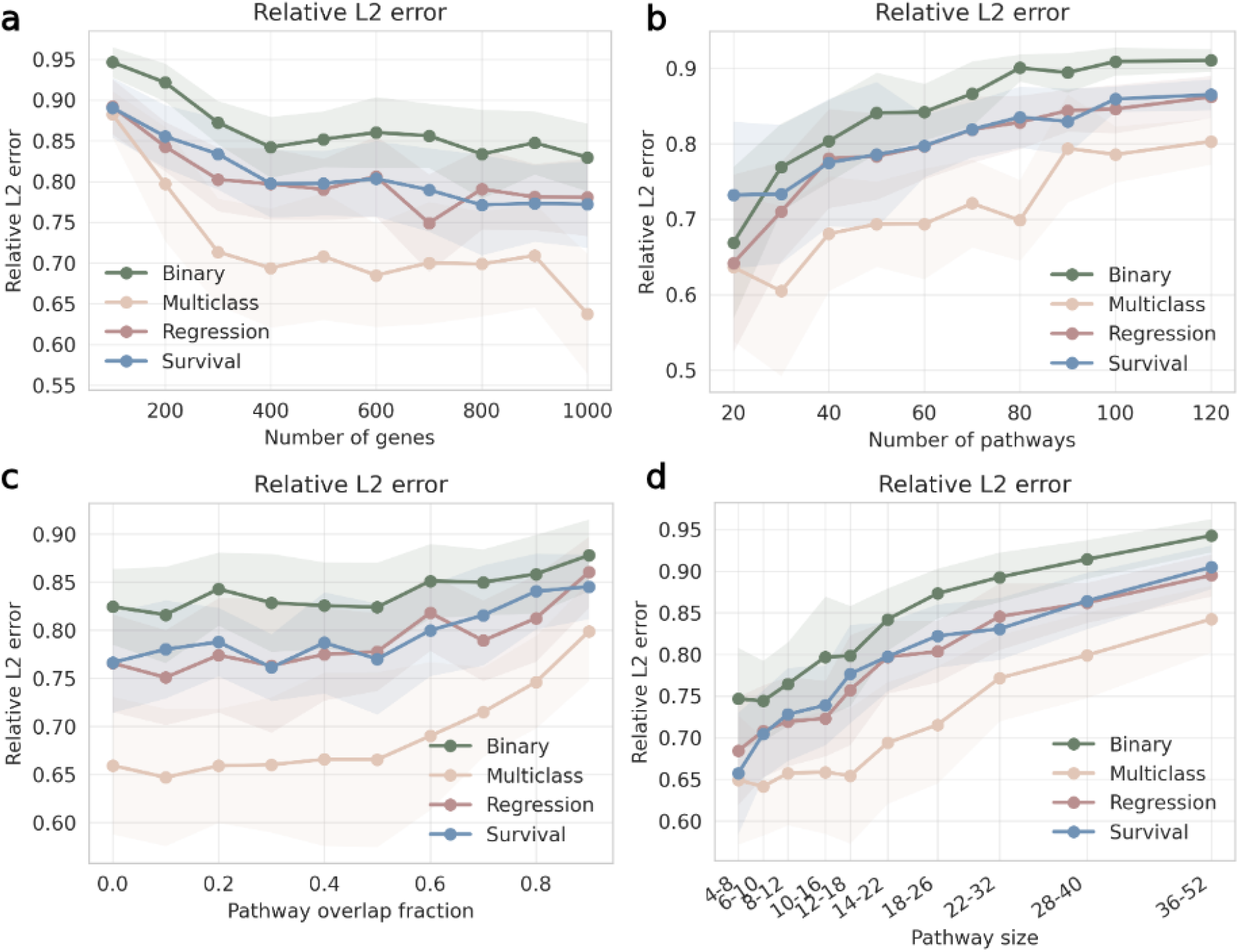
Robustness of nonidentifiability across biological configurations. Relative *L*_2_ error between teacher and student gene-pathway weights under systematic variations of the simulated biological structure. (**a**) Number of genes. (**b**) Number of pathways. (**c**) Pathway overlap fraction. (**d**) Pathway size distribution. Across all configurations and prediction tasks, student networks consistently fail to recover the teacher’s first-layer parameters despite identical architecture and successful output matching. These results demonstrate that the lack of parameter recovery is not tied to a specific choice of simulated biology, but reflects a general property of topology-constrained BINNs. Shaded regions indicate variability across independent runs.

We next examined the role of architectural depth. As shown in Fig. 4, increasing the number of hidden layers progressively degrades recovery of the first biologically interpreted layer. While very shallow networks exhibit slightly lower error, the improvement is modest and recovery remains far from accurate even at minimal depth. Beyond a certain depth, the error increases sharply and saturates close to its maximum value. This behavior is consistent with the increasing ability of deeper networks to compensate for incorrect early representations through downstream transformations, effectively decoupling predictive performance from the structure of the first layer. Importantly, the persistence of poor recovery even in shallow architectures demonstrates that nonidentifiability is not merely a consequence of deep overparameterization, but an intrinsic property of the learning problem.

**Figure 4:**
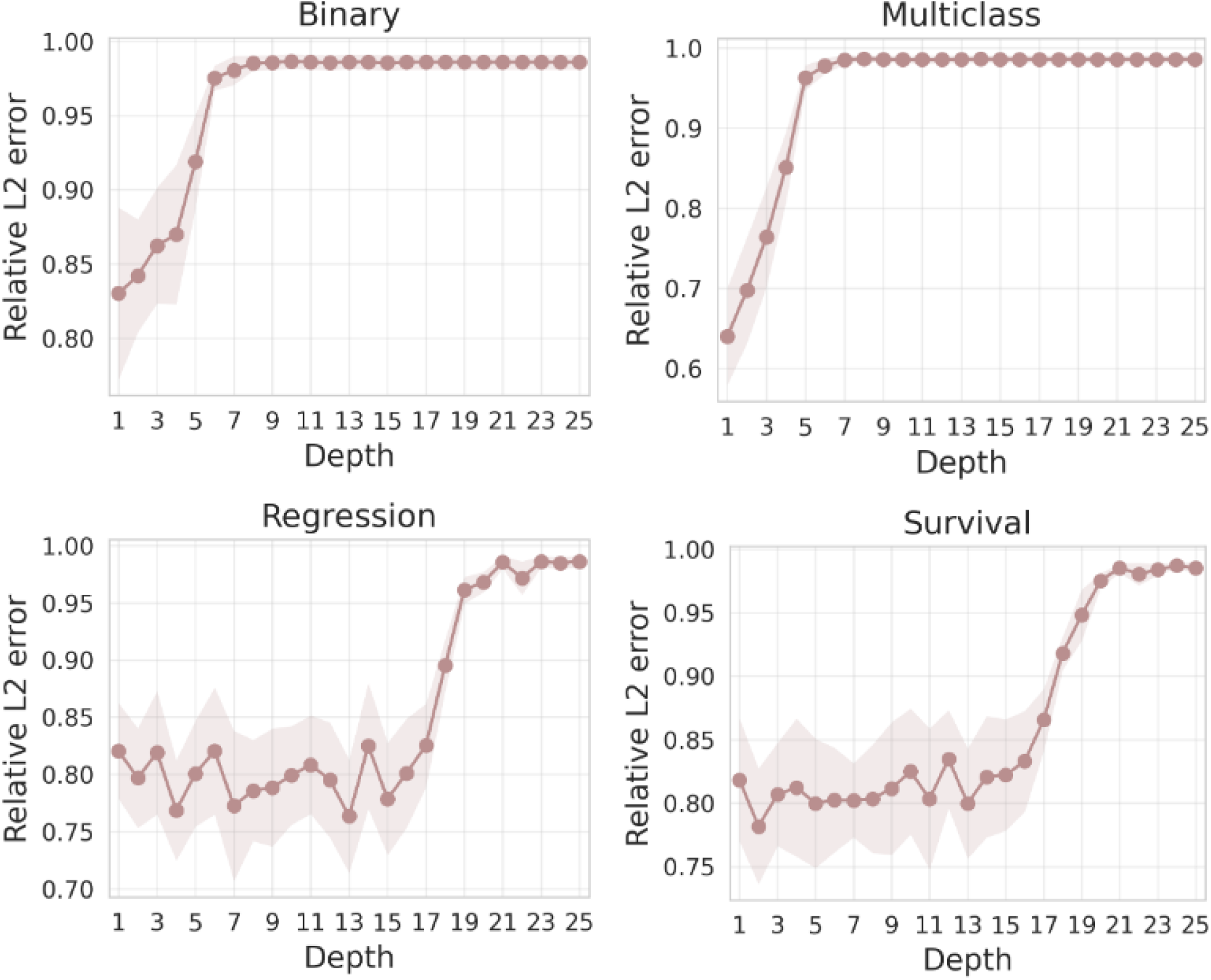
Effect of network depth on parameter recovery. Relative *L*_2_ error between teacher and student gene-pathway weights as a function of the number of hidden layers (**Depth**), across binary classification, multiclass classification, regression, and survival tasks. Increasing network depth progressively degrades recovery of the first biologically interpreted layer, reflecting the growing ability of downstream layers to compensate for incorrect early representations. Notably, although shallower networks exhibit slightly improved alignment, recovery remains consistently poor even at minimal depth, indicating that nonidentifiability is not restricted to deep architectures but is intrinsic to the learning problem. Shaded regions indicate variability across independent runs.

Together, these results indicate that the lack of identifiability is intrinsic to the learning problem and cannot be resolved by architectural choices alone.

## 4 Discussion

Biologically informed neural networks are widely assumed to be interpretable because their architectures reflect known biological structure. Our results challenge this premise directly. Even in a favorable scenario in which the model is correctly specified, the data are noiseless, and the biological wiring is known, these models can recover the correct input-output function while failing to recover both the underlying gene-pathway weights and the corresponding pathway-level activations. Interpretability therefore fails not only at the level of parameters, but also at the level of internal representations.

This failure is not a consequence of imperfect data or modeling choices, but rather a structural property of the learning problem. BINNs are typically trained only to match outputs, such as classification labels, regression targets, or survival endpoints. Because many distinct internal parameterizations can implement the same function, the mapping from data to internal gene-pathway weights is not identifiable in the absence of additional constraints. As a consequence, a model can achieve excellent predictive performance while routing and representing information internally in ways that diverge substantially from the intended biological annotations. Critically, this is not merely a theoretical abstraction. Among the widely adopted BINN implementations reviewed here, interpretability is claimed primarily on the basis of architectural topology, while objective-level constraints on gene-pathway weights or pathway activations are generally absent. The teacher-student framework therefore does not construct an artificial scenario; instead, it models the regime in which these models are commonly used and demonstrates that this regime is insufficient to support the strongest interpretability claims.

This point is well known in machine learning as a problem of nonidentifiability, but it has a specific consequence for biologically informed architectures. In ordinary neural networks, non-identifiability means that internal weights and representations cannot be uniquely recovered from input-to-output observations. In BINNs, the same issue directly affects biological interpretation: a node labelled as a pathway is not necessarily an identified biological variable, and its activation is not necessarily a measurement of pathway activity.

Several mechanisms contribute to this nonidentifiability. Random initialization in high-dimensional parameter spaces is overwhelmingly unlikely to align with biologically meaningful directions, so optimization typically starts far from any biologically interpretable solution. At the same time, neural networks admit large equivalence classes of internal parameterizations: hidden units can be permuted, rescaled, or redistributed across layers without affecting the output [27]. In deeper architectures, this ambiguity is further amplified by the ability of downstream layers to compensate for inaccuracies in earlier representations, effectively decoupling predictive performance from the structure of the biologically interpreted first layer. As a result, the optimization process has no intrinsic preference for solutions that correspond to the underlying biology.

Crucially, this ambiguity extends beyond parameters to the level at which interpretability is often operationalized in practice. Pathway node activations are frequently interpreted as proxies for biological activity, under the assumption that even if weights are not uniquely determined, the learned representations may still be meaningful. Our results show that this assumption is also not guaranteed: networks that produce indistinguishable predictions can rely on substantially different pathway-level activations. Thus, neither gene-pathway weights nor pathway activations can be taken as reliable indicators of biological mechanisms in the absence of additional constraints.

The existence of multiple internal solutions is sometimes loosely interpreted as a form of biological redundancy. However, redundancy explains robustness at the level of function, not identifiability at the level of parameters. The presence of many equivalent internal configurations implies that the mapping from data to internal representations is fundamentally underdetermined. Interpreting any particular configuration as biologically meaningful is therefore not justified without further assumptions.

This underdetermination also clarifies the distinction, introduced above, between topology-based BINNs and models that incorporate prior knowledge through explicit equations or constraints, such as physics-informed neural networks and BioPINN-like approaches. In the latter, biological or physical knowledge enters the optimization objective and restricts the set of admissible solutions. In contrast, in the topology-based BINNs and VNNs considered here, biological knowledge defines the wiring diagram, while interpretability is typically inferred from learned weights or node activations. Our results show that such inference is generally unwarranted.

If mechanistic interpretability is the goal, additional constraints are necessary. The central issue is identifiability: without constraints that break symmetries, restrict the set of equivalent solutions, or anchor internal variables to measured biological quantities, neither parameters nor representations are uniquely determined. Crucially, these constraints must act at the level of the objective function, not only at the level of architecture. To date, such constraints are rarely incorporated systematically in widely used BINNs. The burden of proof, therefore, falls on future work: interpretability cannot be assumed from architecture alone but must be demonstrated through explicit identifiability guarantees.

Incorporating experimentally supported relationships as explicit penalties, fixing or strongly regularizing trusted gene-pathway interactions, or enforcing biologically motivated constraints such as sign or monotonicity can reduce the space of admissible solutions and partially restore identifiability. Standard regularization can also bias the model toward particular solutions [27], but without biologically grounded constraints, it does not guarantee meaningful interpretation.

Taken together, our results expose a fundamental limitation of a widely adopted paradigm. Biologically informed architectures do not, by themselves, guarantee interpretable internal representations. Without explicit constraints that enforce identifiability, both parameters and pathway activations can diverge substantially from the underlying biology while preserving predictive performance. In this sense, the apparent interpretability of these models reflects their design, not what they actually learn.

## Data and Code availability

All code necessary to generate the synthetic data and reproduce the results and figures of this study is available at https://github.com/compbiomed-unito/The-illusion-of-interpretability-in-biologically-informed-neural-networks.

## Additional Information

### Supplementary information for

### S1 Summary of the Supplementary Information

Biological regulation is complex, and in practice we only observe partial and noisy measurements of the underlying mechanisms. In this Supplementary Note we adopt a deliberately simplified setting: we assume that, for the tasks at hand, the relevant biology can be approximated by a neural network whose first (input) layer is *sparse* (e.g., genes as inputs) and whose second layer aggregates inputs into *pathway* nodes (first hidden layer). This abstraction captures the core architectural idea behind biologically-informed neural networks (BINNs): restricting connections according to prior biological knowledge to promote interpretability.

However, even in this idealized regime it is generally *improbable* to recover the “true” gene-to-pathway link values. In other words, architectural sparsity alone does not guarantee weight-level interpretability, and BINNs may not be fundamentally different from standard dense networks in this respect. We formalize this point using a teacher-student paradigm, where the *teacher* represents the (unknown) ground-truth biology and the *student* is the learned BINN.

In noiseless teacher-student experiments, even when the student is trained with the *correct* biological mask, the student can match the teacher’s input-output mapping while still failing to recover the teacher’s first-layer (gene-to-pathway) weights. We provide short demonstrations of two core reasons:

1. **Initialization misalignment:** for a pathway node with *s* incoming genes, the probability that a random isotropic initialization is *positively aligned* (i.e., does not induce an effective sign flip) with a fixed ground-truth weight vector decays exponentially with *s*;
2. **Non-identifiability of first-layer weights:** even under perfect prediction, multiple distinct parameter settings can implement exactly the same function. This can arise from, e.g.,
  - **node permutation** symmetries,
  - **weight rescaling** invariances across layers,
  - **weak first-layer gradients** (many functionally equivalent directions produce negligible changes in loss).

#### Pathway-informed models versus randomized sparse controls

Because sparsity itself can act as an inductive bias, a crucial question is whether *biological* sparsity (curated pathway masks) provides benefits beyond what one would obtain from an arbitrary sparse wiring pattern with the same density and degree distribution. For this reason, alongside pathway-informed BINNs we also consider *randomized sparse controls*: networks whose gene-to-hidden connectivity is randomized while preserving basic sparsity statistics (e.g., number of incoming connections per hidden unit). Comparing these two settings helps disentangle improvements due to genuine biological structure from improvements due merely to reduced parameter count or regularization induced by sparsity.

In the following sections, we provide mathematical justification for the above claims.

### S2 Mathematical analysis

#### S2.1 Setup (masked first layer)

Let *x* ∈ ℝ ^*d*^ denote gene-level inputs. A biologically-informed first hidden layer of size *m* is defined by a binary mask *M* ∈ {0, 1}^m×d^, where *M*_ij_ = 1 indicates gene *j* connects to pathway node *i*. Let *S*_i_ = {*j* : *M*_ij_ = 1} and *s*_i_ = |*S*_i_|.

A teacher network computes

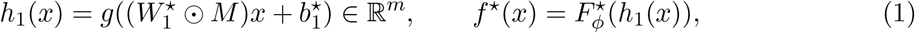

where ⊙ is elementwise multiplication, *g* is applied coordinatewise, and 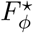 denotes the remaining (typically dense) layers. A student has the same architecture class and the same mask *M*, and is trained (e.g. by backpropagation) to minimize a population loss

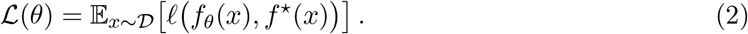

We focus on whether training can recover 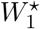

### S2.2 Random initialization almost never aligns with the teacher first-layer weights

#### S2.2.1 Setup (one hidden neuron)

Consider a first hidden neuron *h*_j_ whose input weights form a vector

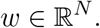

Assume *random initialization* means that the *direction* of *w* is uniformly distributed on the unit sphere

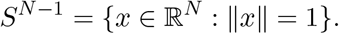

Fix a teacher direction *u* ∈ *S*^*N*−1^. By rotational symmetry we can choose

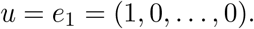

Let *θ* ∈ [0, *π*] be the angle between *w* and *u*, i.e.

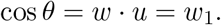

We want the probability that a random direction is within angle *α* of *u*:

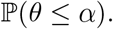

#### S2.2.2 Spherical-cap probability (exact formula)

The event {*θ*≤*α}* is a spherical cap on *S*^*N*−1^. For a uniform point on *S*^*N*−1^, the surface-area density at polar angle *θ* is proportional to sin^*N*−2^ *θ*. Hence the probability of falling in the cap is the ratio of cap area to full sphere area:

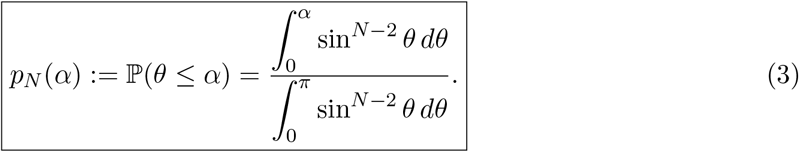

#### S2.2.3 Sanity checks in low dimensions

**Case** *N* = 2. Then sin^*N*−2^ *θ* = sin^0^ *θ* = 1 and

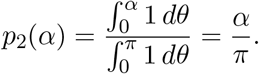

**Case** *N* = 3. Then sin^*N*−2^ *θ* = sin *θ* and

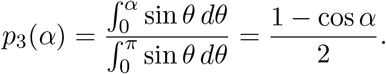

For small *α*, 1 − cos *α* ≃ *α*^2^*/*2, so *p*_3_(*α*) ≃ *α*^2^*/*4.

#### S2.2.4 High-dimensional decay (small-angle scaling)

For small *θ*, sin *θ* ≃ *θ*. Therefore, for small *α*,

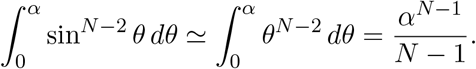

The denominator 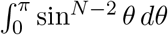 increases exponentially with *N* contributing to the decreasing of *p*_*N*_ (*α*). Hence the dominant dependence on *α* and *N* is

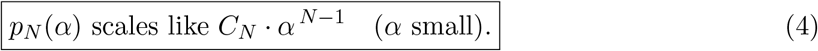

Where for *N* ≥ 2 the factor can be bounded as 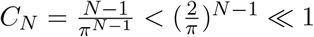. In particular, for any fixed small tolerance angle *α <* 1 (in radians), *p*_N_ (*α*) decreases exponentially fast as *N* increases.

#### S2.2.5 From one neuron to a full first layer of *J* neurons

Let the teacher first layer have *J* input-weight vectors *u*_1_, …, *u*_J_∈ *S*^*N*−1^, and let the student initialize *J* random unit vectors *w*_1_, …, *w*_J_ independently and uniformly on *S*^*N*−1^.

Define *ordered α-alignment* by

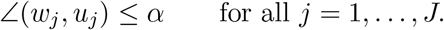

Since the *w*_j_ are independent and each event has probability *p*_*N*_ (*α*),

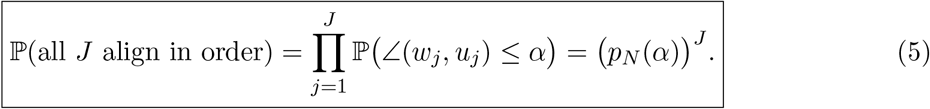

Using the small-angle scaling *p*_*N*_ (*α*) ≈ *C*_*N*_ *α*^*N*−1^, we obtain the dominant behavior

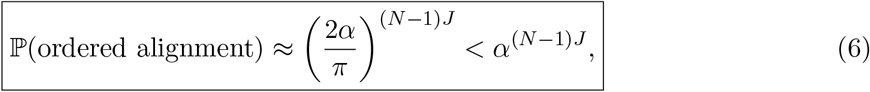

which decreases sharply with both the input dimension *N* and the number of first-layer neurons *J*.

### S2.3 Part 2: perfect function learning does not imply first-layer weight recovery

#### S2.3.1 Teacher-student setting

Let teacher and student share the same feed-forward architecture with *L* ≥ 3 layers and a component-wise nonlinearity *g*(·). The teacher computes

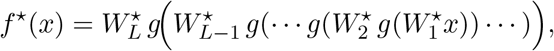

and the student computes *f* (*x*) = *f* (*x*; *W*_1_, …, *W*_L_) with the same form. Training uses an *output-only* loss (e.g. regression/distillation) so that ideally

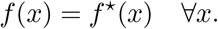

Since the student hypothesis class contains the teacher, zero error is achievable; hence *perfect learning of the mapping is possible*.

#### S2.3.2 Low probability of recovering 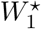 without a specific loss

The key point is *non-identifiability* : if there exists a different parameter set 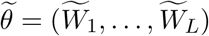 such that

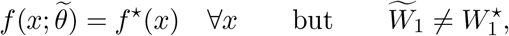

then *no training procedure based only on input-output loss can distinguish θ*^*^ *from* 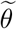 . Indeed, for any loss of the form

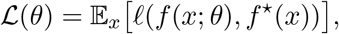

we have

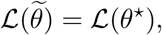

because 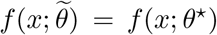 for all *x*. Therefore the set of global minimizers of the output loss typically contains many distinct *θ*, and recovering the teacher’s internal weights is *almost impossible* without additional constraints (e.g. a loss that explicitly matches *W*_1_ or matches internal features).

#### S2.3.3 Explicit source of non-identifiability: permutation symmetry (any component-wise *g*)

For any hidden layer *ℓ* ∈ {1, …, *L* − 2} and any permutation matrix *P*_*ℓ*_ of size *J*_l_ × *J*_l_, define

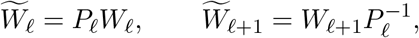

keeping all other layers unchanged. Since *g* is applied component-wise,

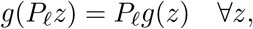

hence the network function is unchanged:

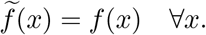

Thus, even after achieving *f* = *f* ^*^, backprop has no preference for the teacher-labeled ordering of hidden units.

Permutation symmetry alone yields at least

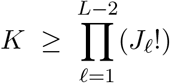

distinct parameterizations implementing the *same* function. Consequently, the probability that training recovers the *exact ordered* teacher parameters (in particular *W*_1_ =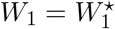) is at most on the order of

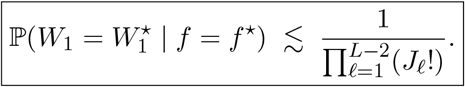

As depth *L* increases, 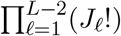 grows extremely fast, so exact reconstruction becomes even less likely.

#### S2.3.4 Stronger (common) effect: continuous non-identifiability (e.g. ReLU)

If *g* is positively homogeneous (e.g. ReLU), then for any diagonal 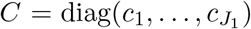 with *c*_j_ > 0,

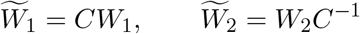

gives the same function. Hence the set of solutions realizing *f* ^*^ typically contains a continuous manifold, and exact recovery of magnitudes in 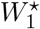 is a measure-zero event:

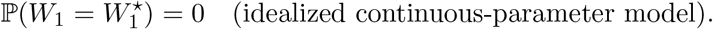

#### S2.3.5 Why depth makes recovery worse in practice: the first-layer gradient shrinks/distorts with *L*

Backpropagation gives

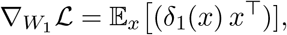

where the error signal *δ*_*ℓ*_ satisfies the recursion

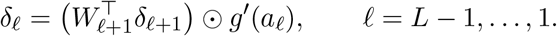

Unrolling,

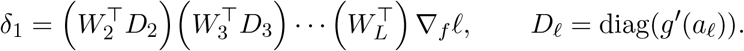

Hence, for any sample *x*,

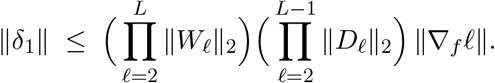

If |*g*^′^(*z*)| ≤ *ρ* (bounded slope; e.g. sigmoid/tanh have *ρ <* 1), then ∥*D*_l_∥_2_ ≤ *ρ* and

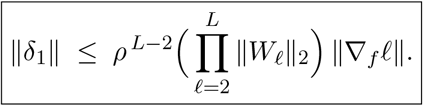

Thus, unless the product of weight norms grows to compensate, the learning signal reaching *W*_1_ decays exponentially in depth. With a sparse mask, the update is further projected by ⊙*M*, reducing effective degrees of freedom and often the update magnitude.

##### Interpretation (matches experiments)

As *L* increases, (i) the equivalence class of parameters implementing *f* ^*^ grows (often continuously), and (ii) the gradient that could move *W*_1_ becomes weaker/more distorted (a product of many Jacobians). Therefore deep students can fit *f* ^*^ by adjusting later layers while leaving *W*_1_ far from 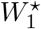 the mismatch typically worsens with depth, consistent with the observed increase of 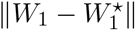 from shallow to deep networks.

#### S2.3.6 Take-home message

Backprop with an output-only loss can learn *f* ^*^ perfectly, but 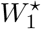 is not identifiable.

The number of functionally equivalent internal representations grows rapidly with depth *L* (at least factorially via permutations, and often continuously via rescalings), so exact first-layer reconstruction without an explicit weight/feature-matching loss is impossible (and in practice, exceedingly unlikely).

### S3 Prediction tasks

We evaluated four prediction tasks commonly encountered in omics-based modeling, spanning classification, regression, and time-to-event analysis. In all cases, the student was trained to reproduce the teacher’s outputs rather than ground-truth labels.

#### Binary classification

The teacher produced binary class probabilities using a sigmoid output layer. The student was trained to match the teacher’s logits using mean squared error loss. Predictive performance was assessed using classification accuracy.

#### Multiclass classification

The teacher generated class probability distributions through a softmax output layer. The student minimized the Kullback-Leibler divergence between teacher and student predictive distributions. Predictive performance was evaluated using classification accuracy.

#### Regression

For regression tasks, the teacher produced continuous scalar outputs through a linear output layer. The student was trained using mean squared error to match teacher predictions. Predictive performance was evaluated using the coefficient of determination *R*^2^ and mean squared error.

#### Survival analysis

For survival tasks, the teacher implemented a Cox proportional hazards model and produced continuous risk scores. The student was trained to match these risk scores using mean squared error. Predictive performance was assessed using the concordance index (C-index), computed on survival times generated by the teacher.

### S4 Evaluation of internal recovery

Internal recovery was evaluated at two complementary levels: parameter recovery and representation recovery.

#### Parameter-level recovery

Recovery of the gene-pathway layer was assessed by comparing corresponding pathway weight vectors between teacher and student networks. For each pathway, we computed the relative *L*_2_ error,

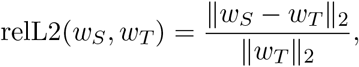

which quantifies deviations in both magnitude and direction, and the cosine similarity,

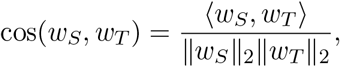

which captures angular alignment independently of scale.

High relative *L*_2_ error or low cosine similarity indicates failure to recover the teacher’s gene-pathway directions, despite successful functional matching at the output level.

#### Representation-level recover

To test whether interpretability might still hold at the level of pathway activity even when weights are not recovered, we also compared teacher and student pathway-node activations for the same input samples. This analysis directly assessed whether pathway nodes in the student encoded the same intermediate representations as the corresponding pathway nodes in the teacher.

Poor activation alignment indicates that pathway nodes do not preserve their intended biological semantics, even when the student accurately reproduces the teacher’s outputs.

Let *z*_T_ (*x*) ∈ ℝ^*P*^ and *z*_S_(*x*) ∈ ℝ^*P*^ denote the pathway-layer activations of the teacher and student networks, respectively, for an input sample *x*, where *P* is the number of pathways. For a given test dataset of *N* samples, we collected the activation matrices *Z*_T_, *Z*_*S*_ ∈ ℝ^*N*×*P*^.

We quantified the similarity between teacher and student activations using three complementary metrics.

First, we computed the Pearson correlation coefficient for each pathway across samples,

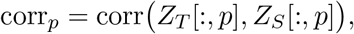

and reported the mean correlation across pathways. This metric captures whether corresponding pathway nodes exhibit consistent activation patterns across samples.

Second, we measured the cosine similarity between activation vectors at the sample level,

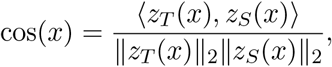

and averaged this quantity across samples. This provides a global measure of alignment between pathway representations for individual inputs.

Third, we computed a global coefficient of determination (*R*^2^) between the two activation matrices,

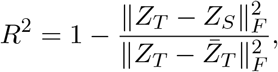

where 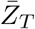 denotes the mean teacher activation across samples. This metric quantifies how well the student reconstructs the overall structure of pathway activations.

All metrics were evaluated on held-out test data and averaged over multiple independent student initializations. Consistently low correlation, cosine similarity, or *R*^2^ values indicate that the student fails to recover the teacher’s pathway-level representations, even in cases where predictive performance is nearly identical.

